# GinJinn2: Object detection and segmentation for ecology and evolution

**DOI:** 10.1101/2021.08.20.457033

**Authors:** Tankred Ott, Ulrich Lautenschlager

## Abstract

1. Proper collection and preparation of empirical data still represent one of the most important, but also expensive steps in ecological and evolutionary/systematic research. Modern machine learning approaches, however, have the potential to automate a variety of tasks, which until recently could only be performed manually. Unfortunately, the application of such methods by researchers outside the field is hampered by technical difficulties, some of which, we believe, can be avoided.
2. Here, we present GinJinn2, a user-friendly toolbox for deep learning-based object detection and instance segmentation on image data. Besides providing a convenient command-line interface to existing software libraries, it comprises several additional tools for data handling, pre- and postprocessing, and building advanced analysis pipelines.
3. We demonstrate the application of GinJinn2 for biological purposes using four exemplary analyses, namely the evaluation of seed mixtures, detection of insects on glue traps, segmentation of stomata, and extraction of leaf silhouettes from herbarium specimens.
4. GinJinn2 will enable users with a primary background in biology to apply deep learning-based methods for object detection and segmentation in order to automate feature extraction from image data.

## Introduction

Conducting empirical studies in ecology and evolutionary/systematic biology typically requires substantial effort for proper data collection and preparation. The ability to automate time-consuming steps of this process, possibly along with further downstream analyses, for example, using programming languages like Python or R, can not only increase productivity, but also allow otherwise infeasible large-scale analyses. Recent advances in machine learning (ML), both on the soft- and hardware side, make it even possible to automate tasks that are difficult to solve by means of classically designed algorithms. Computer vision, in particular, has largely profited from deep learning, which increasingly influences even the more traditional branches of organismic biology. Species identification tools running on smartphone devices (for an overview, see Wäldchen & Mäder, 2018; Jones, 2020) are prominent examples for this trend. Beyond pure classification tasks, a technically even more challenging problem consists in localizing objects like cells, organs, or individuals on images. Specialized tools address this problem for various areas of application, such as crop or weed detection (e.g., Buddha et al., 2019; Afonso et al., 2020), detection of leaves and other plant organs on herbarium specimens (e.g., Ott et al., 2020; Weaver et al., 2020; Younis et al., 2020), stomata counting using microscopic leaf images (e.g., Fetter et al., 2019), animal counting using camera traps (Norouzzadeh et al., 2021), and many more.

Despite the availability of increasingly convenient frameworks, adapting well-established ML methods to new areas of application typically requires an amount of technical knowledge that may discourage potential users. GinJinn2, whose core functionality is based on Detectron2 (Wu et al., 2019), aims at lowering this hurdle by providing an easy-to-use command-line interface to the latter, augmented by a number of utility functions, designed to help the user with building custom analysis pipelines. While its predecessor (Ott et al., 2020) focussed on extracting leaves from digitized herbarium specimens, the presented program is a complete reimplementation, aiming at a wider scope of application. Unlike the former, it is not restricted to bounding-box object detection, but also incorporates functionality for instance segmentation, i.e., pixel-precise detection and classification of individual objects.

In the present contribution, a number of example analyses demonstrate how ecological, agricultural or evolutionary/systematic studies may benefit from GinJinn2. Those include pest monitoring using yellow glue traps, leaf-shape extraction from herbarium specimens, stomata segmentation, and the evaluation of seed mixtures. We hope to encourage interested researchers to consider deep learning-based object detection or segmentation when faced with similar tasks. Using GinJinn2 together with pretrained models from Detectron2’s model zoo, new applications can be explored with a minimum of invested time and effort, which makes it a potentially useful tool for both beginners and advanced users.

## Software

### Overview

GinJinn2 is a toolbox for deep learning-based bounding-box object detection and instance segmentation. As such, it provides functionality for model training, evaluation and application based on the Detectron2 framework, segmentation refinement based on CascadePSP (Cheng et al., 2020), a set of data pre- and postprocessing tools for handling annotated image datasets, and capabilities for data insight and visualization. GinJinn2 is not meant to be a replacement for existing frameworks like Detectron2 or the Tensorflow Object Detection API (Huang et al., 2017), but rather a toolkit enabling code-free access to deep learning-based object detection technologies. All of GinJinn2’s functionality is accessible via an easy-to-use command-line interface (CLI).

### Dataset splitting

Before training one of the available object-detection models, the user may want to split annotated image data into two or three sub-datasets. Besides the data used to train the model, it is generally advisable to use a so-called validation dataset in order to detect overfitting and to optimize model choice and training parameters. Using a separate dataset for those purposes is necessary because the model’s fit to the training data does not provide information about its generalization capability. In other words, a trained model may accurately reproduce the training data, but perform poorly on images that have not been presented to it before. However, as soon as any optimizing decision has been made based on the validation data (e.g., when to stop the training process), the model may again show overly optimistic performance for this particular dataset. To obtain an unbiased evaluation of the final model, it is therefore necessary to provide an additional test dataset, which should not have been used for any other task beforehand. The *ginjinn split* command partitions an input dataset in such a way that each image along with its annotated objects is assigned to one of the resulting subsets. To be representative for the original dataset, each of the latter should comprise similar proportions of objects from each category. Aiming at a high level of homogeneity, the proposed splits are generated by a greedy optimization algorithm. Despite being a relatively rough heuristic, this approach is often sufficient to create acceptable splits and can even be applied to large datasets.

### Object detection and instance segmentation

GinJinn2, by leveraging Detectron2’s model zoo, offers several Faster R-CNN (Ren et al., 2015) and Mask R-CNN (He et al., 2017) models for bounding-box detection and instance segmentation, respectively. These are used in a supervised manner, i.e., before being able to predict objects on new images in a meaningful way, their parameters (“weights”) have to be fitted to images with known object occurrences (“training”). While training such models *de novo* can be highly GPU-intensive, this process can be considerably abbreviated by starting from pretrained rather than randomly initialized weights (“transfer learning”). Accordingly, all available Detectron2 models have already been trained on a large image dataset. Using those pretrained networks reduces the training time for new, custom datasets as well.

Once the user has prepared datasets for training, and, optionally, validation and test (see Dataset splitting), a GinJinn2 project can be initialized using *ginjinn new*. Training models using *ginjinn train* constitutes the computationally most demanding part of a typical GinJinn2 pipeline. This process consists of a prespecified number of iterations, at each of which one or multiple images from the training dataset are presented to the model. The objects predicted by the latter are then compared to the known annotations and the model weights are adjusted to reduce deviations (“loss”) from the desired output. While minimizing the loss with respect to the training dataset, at some point, the model’s generalization capability may begin to degrade. This so-called overfitting can be recognized by an increasing loss for the validation dataset. The latter is therefore evaluated at predefined intervals. To enable a better assessment of the learning progress, COCO (Lin et al., 2014) evaluation metrics for the validation dataset are calculated as well. Since the model weights are stored periodically, in case of overfitting, the user can go back to an earlier checkpoint without having to discard the complete training. Since GinJinn2 is using Detectron2 as modelling backend, all models that are trained in the context of a GinJinn2 project can be used with Detectron2’s Python interface without modification.

The quality of the final, trained model is best assessed based on a hitherto unused dataset with known object occurrences. This can be done using *ginjinn evaluate*, which calculates COCO evaluation metrics for the specified test dataset.

The *ginjinn predict* command allows applying a trained model to predict object occurrences for arbitrary images. Instance segmentations can optionally be refined using CascadePSP (Cheng et al., 2020); while slowing down the predictions, this may considerably improve the quality of the object outlines, especially in case of clear object boundaries. To facilitate the further use of the predictions, GinJinn2 provides various output options: 1) visualization of the predictions on the original images, 2) writing a new COCO annotation file, and 3) saving a cropped image and, if applicable, segmentation mask for each predicted object.

### Further functionality

#### GinJinn2 offers several utilities for data pre- and postprocessing

As a counterpart to the already described splitting command (*ginjinn split*), datasets can also be merged (*ginjinn utils merge*), which is particularly useful when using COCO’s annotation format. In doing so, the input datasets are also checked for duplicated images.

Object annotations can be filtered by either category or size using *ginjinn utils filter_cat* or *ginjinn utils filter_size*, respectively. The latter command is also capable of removing only small disjunct fragments from existing objects.

To simplify existing data, nested image directories can be summarized, making them compatible with GinJinn2 and other tools. *ginjinn utils flatten* recursively collects all images from a given directory and its sub-directories, renames and copies them into a single directory, and modifies associated annotations accordingly.

Due to the limited spatial resolution of common object detection models, detecting or segmenting objects that are small in relation to the image size can be difficult. To mitigate this problem, a sliding-window approach can be used to split the original images into smaller sub-images (*ginjinn utils sw_split*), preserving annotated objects, if available. Conversely, predictions based on such fragmented images can be merged again (*ginjinn utils sw_merge*) in order to generate an annotation of the original image.

The *ginjinn utils crop* command creates an annotated sub-image for each annotated object from a given dataset. Similar to the sliding-window approach, this can be utilized to increase objects sizes relative to the images. Specifically, performing instance segmentation based on previously cropped bounding boxes may lead to improved results.

#### Besides the aforementioned data processing features, the following commands aim to provide additional convenience

The contents of a dataset can be briefly summarized using *ginjinn info*. More detailed information is provided by *ginjinn utils count*, which lists object occurrences individually for each image in a given dataset. Object annotations can be visualized with *ginjinn visualize*, which produces images overlaid by bounding boxes and, if available, segmentation polygons. Moreover, Ginjinn2 allows to generate artificial datasets for testing purposes (*ginjinn simulate*).

### Installation and usage

GinJinn2 is implemented in Python3 and can be installed using the Conda package manager, which also takes care of most of its dependencies. *ginjinn* and all its subcommands provide a help option to list available parameters along with a short description. Further guidelines regarding installation and usage, along with an introductory tutorial and exemplary applications, are provided at https://ginjinn2.readthedocs.io.

## Example analyses

### Seed counting

In this section, we demonstrate how GinJinn2 can be applied for seed mixture analysis, an illustrative use case for bounding-box detection with subsequent counting. This approach could, for instance, be used to examine commercial seed mixtures or be applied to ecological samples (e.g., from seed traps). The presented analysis is based on a dataset consisting of 284 microscopic images of sand-contaminated seed mixtures of the two plant genera *Sedum* L. and *Arabidopsis* (DC.) Heynh.

For all images, intact seeds were annotated with bounding boxes using the Computer Vision Annotation Tool (CVAT, https://github.com/openvinotoolkit/cvat), resulting in 6,732 and 1,964 annotated seeds for *Arabidopsis* and *Sedum*, respectively. The annotated images were exported as COCO dataset, which was then flattened (*ginjinn utils flatten*), and split into sub-datasets for training, validation, and testing. A Faster R-CNN model was simultaneously trained and validated (Figure 1A). The quality of the fit model was assessed using COCO evaluation metrics for bounding-box detection. In addition, instances predicted for the test dataset were counted (*ginjinn utils count*) and compared with the manually obtained counts.

**Figure 1.**
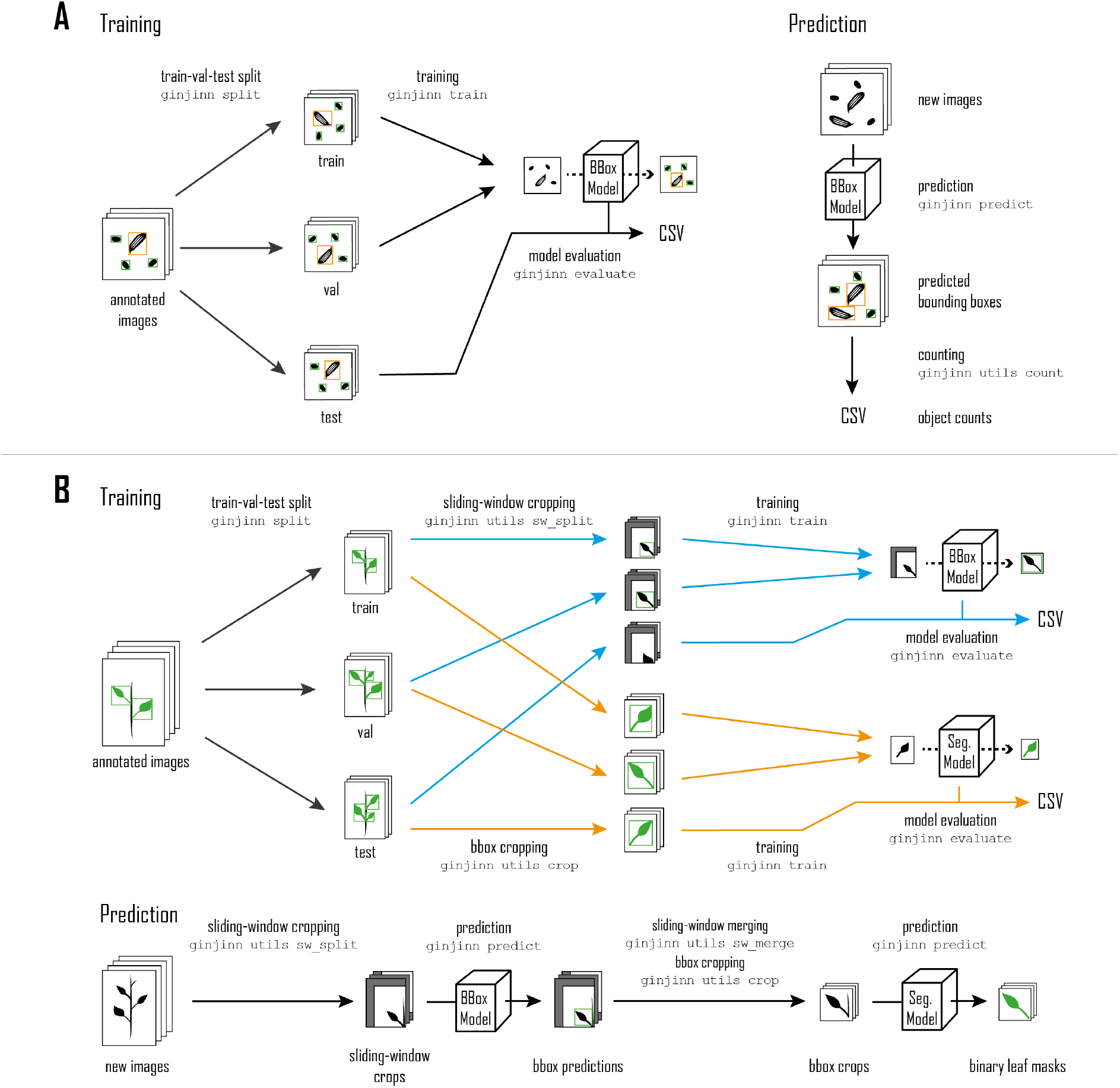
Seeds (**A**) and *Leucanthemum* (**B**) analysis workflows. The Seeds dataset is split into training, validation, and test datasets, which are used to train and evaluate a bounding-box model (**A**, Training). The trained model is applied to new data for seed counting (**A**, Prediction). The *Leucanthemum* dataset is also split into training, validation, and test datasets, but the workflow comprises training and evaluation of two separate models (**B**, Training). The blue branch refers to a bounding-box model for the detection of leaves on sliding-window crops of the split dataset. The orange branch depicts the training and evaluation of an instance segmentation model on padded bounding boxes cropped from the split datasets. Leaf segmentations for new data are predicted by combining both models (**B**, Prediction).

After training, the AP50 was 98.6 and 98.90 for the validation and test dataset, respectively, which indicates that no overfitting occurred. The mean absolute error (MAE) of the class counts for the training dataset was 0.77 for *Arabidopsis* and 0.58 for *Sedum*, meaning that on average, less than a single object per image was misclassified, missed, or falsely detected. The MAE of the seed proportions was 0.01, i.e., only one percent deviation from the true seed proportions.

### Yellow-sticky-traps insect detection and counting

As an example project for counting small, low-contrast objects on large images, the Yellow-Sticky-Traps dataset (Nieuwenhuizen, 2018) was analyzed. This dataset consists of images of yellow glue traps that were placed in greenhouses to monitor insect abundance. Three categories of insects (true bugs) were annotated with bounding boxes: Whitefly (WF), *Macrolophus* (MR), and *Nesidiocoris* (NC).

After removing redundant images and correcting erroneous or missing annotations using CVAT, a cleaned sub-dataset comprising 120 images along with 4,913 bounding-box annotations (WF: 3,660, MR: 1,069, NC: 184) was exported in COCO format. In contrast to the seeds dataset, these bounding-box annotations are of considerably lower quality, often enclosing the insects only loosely.

The cleaned dataset was split into training, validation, and test datasets using *ginjinn split*. Since the insects are relatively small compared to the total image size, a sliding-window approach was applied (*ginjinn utils sw_split*) to crop sub-images along with corresponding object (sub-)annotations. The cropped datasets were used to train and evaluate a Faster R-CNN model for bounding-box detection. Finally, object instances predicted for the test dataset were counted (*ginjinn untils count*) and compared with true object counts.

The trained model achieved a validation and test AP50 of 90.12 and 92.4, respectively. The mean absolute error (MAE) of the instance counts was 1.67 for WF, 0.21 for NC, and 0.79 for MR at an average of 27.1, 1.67, and 7.41 annotated instances per image for the respective object categories. The former amounts to a relative counting error of 6% for WF, 12.5% for NC, and 10.6% for MR (weighted average: 7.24%).

### Stomata segmentation

To demonstrate basic instance segmentation with the aim of detecting stomata, we applied GinJinn2 to microscopic images of epidermal plant material, retrieved from the Cuticle Database Project (Barclay et al., 2012). Results of such a segmentation can be used in downstream analyses for counting, measuring density, or examining size and shape of the stomata.

Using CVAT, 147 images were annotated with 2,314 polygons, each enclosing the guard cells of a stoma. The annotated images were exported as COCO dataset and split into training, validation, and test datasets used to train and evaluate a Mask R-CNN model.

The trained model achieved an AP of 49.46 and 51.32 for the validation and test dataset, respectively. The mean absolute counting error amounts to 2.34 at an average of 14.69 stomata per image. An exemplary prediction is shown in Figure 2A.

**Figure 2.**
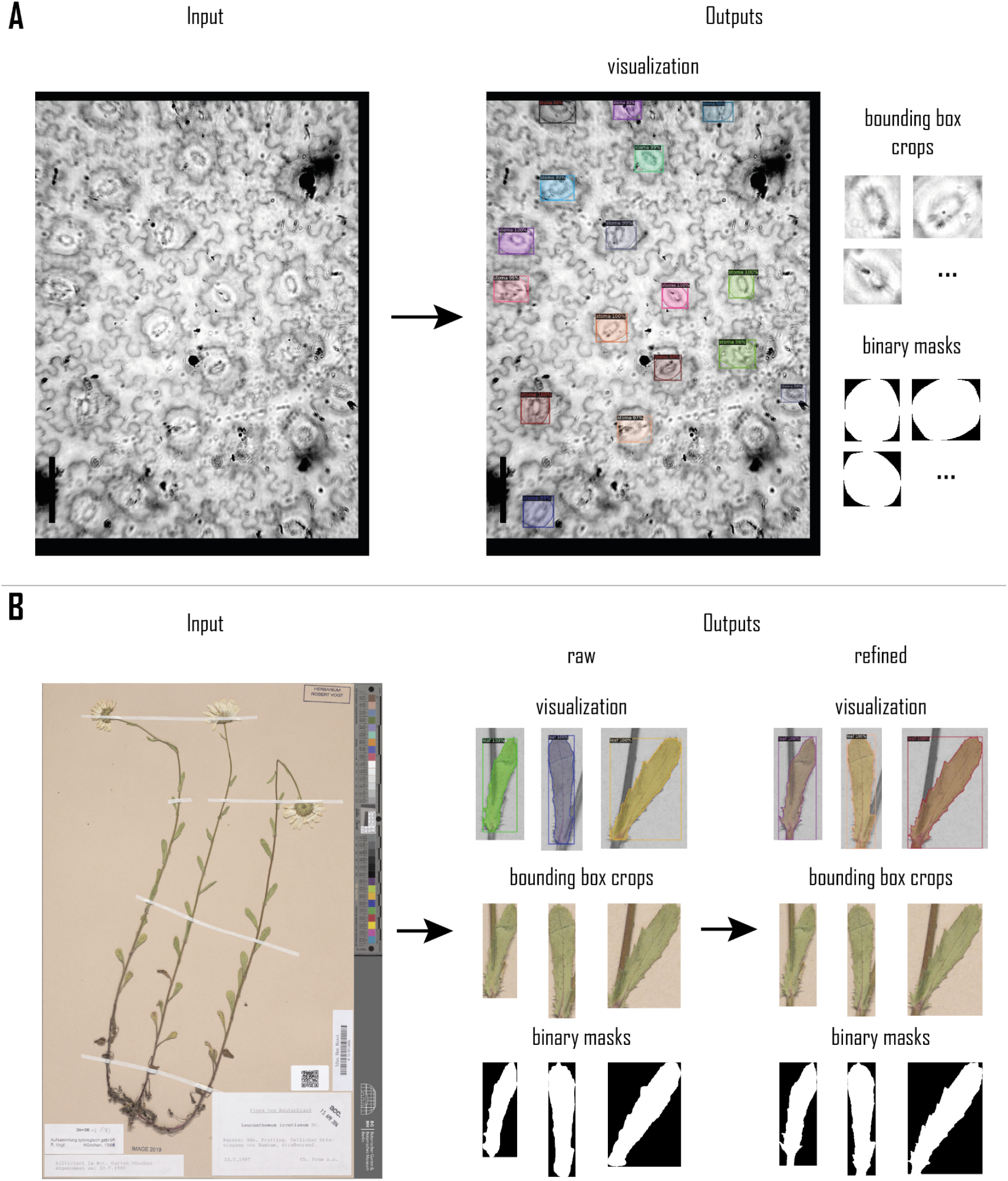
Exemplary outputs from the Stomata (**A**) and *Leucanthemum* (**B**) analyses. **A** depicts a single input image along with corresponding predictions by the stomata model, showing different output formats. Similarly, **B** shows an input image and corresponding predictions for the *Leucanthemum* pipeline, before and after segmentation refinement.

### *Leucanthemum* leaf segmentation

Morphometric studies often rely on outline data of specific animal or plant organs like, for example, leaves in the latter organism group. A common workflow to generate such data is to manually remove leaves from a living or herborized plant, fixate them on a contrasting surface, capture digital images, and finally apply semi-automatic thresholding methods (e.g., OTSU-thresholding) to construct binary segmentation masks. In this exemplary application of GinJinn2, we show an alternative way to segment individual leaves from digitized herbarium specimens based on a two-step approach involving separate models for bounding-box detection and segmentation.

For this purpose, the Botanic Garden and Botanical Museum Berlin provided us with 303 digitized herbarium specimens from 12 different *Leucanthemum* Mill. (ox-eye daisy) species. Using CVAT, the specimen images were annotated with polygons of the single object category “leaf”. This category represents largely intact leaves, which are a prerequisite for reliable morphometric analyses. The annotated images, comprising 950 “leaf” instances, were exported from CVAT as COCO dataset, flattened (*ginjinn utils flatten*) and split into training, validation, and test datasets.

A two-step pipeline (Figure 1B) was applied, consisting of 1) a Faster R-CNN bounding-box detection model that allows to extract individual leaves, and 2) a Mask R-CNN model to segment the leaves on those image parts. The Faster R-CNN was trained and evaluated on sliding-window crops (*ginjinn utils sw_split*) of the three datasets. For the Mask R-CNN, sub-images (*ginjinn utils crop)* were cropped from the original annotated images, each containing a single annotated leaf. Based on those cropped datasets, the Mask R-CNN was trained and evaluated. In addition, segmentation refinement was applied to the predictions for the test dataset.

After training, the Faster R-CNN achieved an AP of 30.57 and 25.85 for the validation and test dataset, respectively. The Mask R-CNN’s AP scores were 76.44 and 74.54. Figure 2B illustrates an exemplary prediction. For new image data, the complete prediction process also involves sliding-window merging as illustrated in Figure 1B in order to remove duplicated objects.

All used GinJinn2 commands and the corresponding project configuration files can be found in the Supporting Information (S1-S6).

## Discussion

The GinJinn2 framework advances the original GinJinn by reimplementing its ideas on the basis of Detectron2, while also introducing new features like segmentation models including mask refinement, as well as several data pre- and postprocessing capabilities.

Based on four exemplary datasets we have shown applications of varying complexity. The seeds and yellow-sticky-traps analyses address multi-category object counting problems using bounding-box detection. We were able to predict the seed ratios with an absolute error of only 1%, proving the potential of our software for the automation of such counting tasks. Considering the similar problem of counting insects on yellow glue traps, with an error of 7.2%, the accuracy of the trained model may appear less convincing. There are two likely causes for this difference in accuracy: 1) low contrast between objects (insects) and background (glue trap) and 2) low quality of annotations. The latter could easily be solved by a more careful annotation scheme. Nevertheless, the achieved accuracy might be sufficient for practical applications, e.g., to measure the response to insecticide treatments or released beneficials in greenhouses.

The stomata analysis serves as a basic example of instance segmentation. Despite several previous works on the automated examination of stomata (Toda et al., 2018; Fetter et al., 2019; Li et al., 2019; Carrasco et al., 2020; Casado-García et al., 2020; Meeus et al., 2020; Song et al., 2020), this contribution, to our knowledge, is the first trying to automatically segment whole stomata (represented by their guard cells) using deep learning. With only 88 highly variable training images, our model achieved an AP of 51.32. Depending on the intended downstream analyses, this precision may already be acceptable if, for instance, only few high-quality object instances are required. Undoubtedly, a model trained on a larger dataset will achieve substantially higher predictive power.

Finally, the *Leucanthemum* analysis illustrates how to construct a pipeline consisting of sliding window-based bounding-box detection and subsequent segmentation for the extraction of high-quality leaf silhouettes from herbarium specimens. Here, the Faster R-CNN achieved an AP of 25.85. For potential morphometric analyses, we are not interested in extracting all leaves, but only largely intact ones, even at the cost of discarding viable instances. Therefore, the relatively low AP is sufficient. The Mask R-CNN, with an AP of 74.54 before refinement, was very successful at segmenting the leaves inside the bounding boxes. This pipeline already allows to generate leaf outlines for downstream analyses like Elliptic Fourier Analysis or Leaf Dissection Index calculation (for an overview of such methods, see McLellan & Endler, 1998) with little manual effort.

With the presented exemplary analyses, we hope to provide guidance for the application of GinJinn2 for automatic data collection and feature extraction. Despite GinJinn2’s progress compared to its predecessor, there is still room for further improvements. At the moment, GinJinn2 is only available for Unix-like operating systems with access to an NVidia GPU while Windows support may become available with forthcoming updates to the Windows Subsystem for Linux (WSL). Moreover, there is only one meta-architecture for each of the two detection tasks available, namely Faster R-CNN and Mask R-CNN. These, however, are among the most successful architectures for general-purpose object detection and segmentation. The integration of additional model architectures may be part of future versions.

We are confident that GinJinn2 will enable users, even those without programming experience, to apply deep learning-based methods for object detection and segmentation as part of their analysis pipelines. Besides, advanced users may utilize GinJinn2 as a tool for rapid prototyping.

## Acknowledgements

First and foremost, we would like to thank Christoph Oberprieler (Regensburg), who acquired the funding, for enabling this work and his comments on the manuscript. We thank Robert Vogt (Berlin) and Sergey Rosbakh (Regensburg) for providing digital images of *Leucanthemum* specimens and seed mixtures, respectively. We would also like to thank David Dilcher (Bloomington) for granting us permission to use the microscopic image shown in Figure 2A. The help of Maximilian Schall and Sebastian Segieth (both Regensburg), who annotated many of the *Leucanthemum* and seeds images, is much appreciated. We thank Tanja Wenzel for her support in designing the workflow diagrams. We also thank Agnes Scheunert (Regensburg) for her comments on the manuscript.

This work was supported by a Grant (OB 155/13-1) of the German Research Foundation (DFG) in the frame of the Priority Programme SPP 1991 “Taxon-omics – New Approaches for Discovering and Naming Biodiversity” to Christoph Oberprieler).

## Author contributions

TO and UL envisioned the present work, implemented the software, carried out the analyses, and wrote the manuscript. Both authors approved the final version of the manuscript. We further note that UL and TO contributed equally to this work. The order of their names in the author list was decided by coin toss.

## Data availability

GinJinn2’s source code and manual are freely available at GitHub (https://github.com/AGOberprieler/GinJinn2). The annotated Seeds, Yellow-sticky-traps and *Leucanthemum* datasets are hosted by the German Federation for Biological Data (GfBio; Link A, Link B, Link C; will be supplied as soon as available). The images used for the Stomata analysis are hosted by the Cuticle Database (Barclay et al., 2012), a Python script for splitting the images is provided in the supporting information (S7); the corresponding annotations are hosted by GfBio (Link D; will be supplied as soon as available).

## Supporting information

commands.pdf:

Appendix S1. GinJinn2 commands of exemplary analyses. seeds.yaml:

Appendix S2. GinJinn2 configuration file for the seeds analysis. stickytraps.yaml:

Appendix S3. GinJinn2 configuration file for the yellow-sticky-traps analysis. stomata.yaml:

Appendix S4. GinJinn2 configuration file for the stomata analysis. leucanthemum_bbox.yaml:

Appendix S5. GinJinn2 configuration file for the *Leucanthmum* analysis (bounding-box detection).

leucanthemum_segmentation.yaml:

Appendix S6. GinJinn2 configuration file for the *Leucanthmum* analysis (instance segmentation).

split_image.py:

Appendix S7. Image splitting script.

## Notes

### Competing Interest Statement

The authors have declared no competing interest.

