## Supplementary material for "GinJinn2: Object detection and segmentation for ecology and evolution": commands.pdf

### Seeds

1. train-validation-test split

   ```
   ginjinn split \
       -I seeds \
       -o seeds_split \
       -d bbox-detection
   ```

2. generate new GinJinn2 project

   ```
   ginjinn new seeds_split_analysis \
       -d seeds_split \
       -t faster_rcnn_R_101_FPN_3x.yaml \
   ```

3. edit seeds_split_analysis/ginjinn_config.yaml
4. train model

   ```
   ginjinn train seeds_split_analysis
   ```

5. evaluate model

   ```
   ginjinn evaluate seeds_split_analysis
   ```

6. (optional) predict on test dataset (or new images)

   ```
   ginjinn predict seeds_split_analysis \
       -i seeds_split/test/images
       -o seeds_split_test_prediction \
       -c -v
   ```

### Stickytraps

1. train-validation-test split

   ```
   ginjinn split \
       -I stickytraps \
       -o stickytraps_split \
       -d bbox-detection
   ```

2. sliding-window cropping

   ```
   ginjinn utils sw_split \
       -I stickytraps_split \
       -o stickytraps_split_sw \
       -s 1024 -p 256 -c
   ```

3. generate new GinJinn2 project

   ```
   ginjinn new stickytraps_split_sw_analysis \
       -d stickytraps_split_sw \
       -t faster_rcnn_R_101_FPN_3x.yaml \
   ```

4. edit stickytraps_split_sw_analysis/ginjinn_config.yaml

5. train model

   ```
   ginjinn train stickytraps_split_sw_analysis
   ```

6. evaluate model ginjinn evaluate stickytraps_split_sw_analysis

7. (optional) predict on test images (or new images)

   7.1 sliding-window crop images

   ```
   ginjinn utils sw_split \
       -i stickytraps_split_sw/test/images/ \
       -o sticktraps_newdata_sw \
       -s 1024 -p 256
   ```

   7.2 detect bounding boxes

   ```
   ginjinn predict stickytraps_split_sw_analysis \
       -i sticktraps_newdata_sw \
       -o sticktraps_newdata_sw_prediction \
       -c -v
   ```

   7.3 (optional) merge sliding-window predictions to reconstruct predictions on original images

   ```
   ginjinn utils sw_merge \
       -a sticktraps_newdata_sw_prediction/annotations.json \
       -i sticktraps_newdata_sw \
       -o sticktraps_newdata_sw_prediction_merged \
       -t bbox-detection
   ```

   7.4 (optional) visualization of the merged predictions

   ```
   ginjinn vis \
       -I sticktraps_newdata_sw_prediction_merged \
       -v bbox
   ```

### Stomata

1. train-validation-test split

   ```
   ginjinn split \
       -I stomata \
       -o stomata_split \
       -d instance-segmentation
   ```

2. generate new GinJinn2 project

   ```
   ginjinn new stomata_split_analysis \
       -d stomata_split \
       -t mask_rcnn_R_101_FPN_3x.yaml \
   ```

3. edit stomata_split_analysis/ginjinn_config.yaml
4. train model

```
ginjinn train stomata_split_analysis
```

5. evaluate model

```
ginjinn evaluate stomata_split_analysis
```

6. (optional) predict on test images (or new images)

```
ginjinn predict stomata_split_analysis \
    -i stomata_split/test/images \
    -o stomata_prediction \
    -c -v
```

### Leucanthemum

1. flatten CVAT output

```
ginjinn utils flatten \
    -i leucanthemum/images/ \
    -a leucanthemum/annotations/instances_default.json \
    -o leucanthemum_flat
```

2. remove empty categories

```
ginjinn info -I leucanthemum_flat
ginjinn utils filter_cat \
    -a leucanthemum_flat/annotations.json \
    -i leucanthemum_flat/images/ \
    -o leucanthemum_flat_filtered \
    -d -f main_vein
ginjinn info -I leucanthemum_flat_filtered
```

3. train-validation-test split

```
ginjinn split \
    -I leucanthemum_flat_filtered \
    -o leucanthemum_flat_filtered_split \
    -d instance-segmentation
```

4. bounding-box model

   4.1 sliding-window cropping

```
ginjinn utils sw_split \
    -I leucanthemum_flat_filtered_split \
    -o leucanthemum_flat_filtered_split_sw \
    -s 2056 -p 512 -c
```

   4.2 generate new GinJinn2 project

```
ginjinn new leucanthemum_flat_filtered_split_sw_analysis \
    -t faster_rcnn_R_101_FPN_3x.yaml \
    -d leucanthemum_flat_filtered_split_sw/ \
```

4.3 edit leucanthemum_flat_filtered_split_sw_analysis/ginjinn_config.yaml

4.4 train model

```
ginjinn train leucanthemum_flat_filtered_split_sw_analysis
```

4.5 evaluate model

```
ginjinn evaluate leucanthemum_flat_filtered_split_sw_analysis
```

4.6 (optional) predict on test dataset

```
ginjinn predict leucanthemum_flat_filtered_split_sw_analysis \
    -i leucanthemum_flat_filtered_split_sw/test/images \
    -o leucanthemum_flat_filtered_split_sw_test_prediction \
    -c -v
```

4.7 (optional) merge sliding-window predictions to reconstruct predictions on original images

```
ginjinn utils sw_merge \
    -a leucanthemum_flat_filtered_split_sw_test_prediction/annotations.json \
    -i leucanthemum_flat_filtered_split_sw/test/images \
    -o leucanthemum_flat_filtered_split_sw_test_prediction_merged \
    -t bbox-detection
```

4.8 (optional) visualization of the merged predictions

```
ginjinn vis \
    -I leucanthemum_flat_filtered_split_sw_test_prediction_merged \
    -v bbox
```

5. segmentation model

5.1 bounding box cropping

```
ginjinn utils crop \
    -I leucanthemum_flat_filtered_split \
    -o leucanthemum_flat_filtered_split_cropped \
    -t segmentation \
    -p 25
```

5.2 generate new GinJinn2 project

```
ginjinn new leucanthemum_flat_filtered_split_cropped_analysis \
    -t mask_rcnn_R_101_FPN_3x.yaml \
    -d leucanthemum_flat_filtered_split_cropped
```

5.3 edit leucanthemum_flat_filtered_split_cropped_analysis/ginjinn_config.yaml

5.4 train model

```
ginjinn train leucanthemum_flat_filtered_split_cropped_analysis
```

5.5 evaluate model

```
ginjinn evaluate leucanthemum_flat_filtered_split_cropped_analysis
```

5.6 (optional) predict on test dataset

```
ginjinn predict leucanthemum_flat_filtered_split_cropped_analysis \
    -i leucanthemum_flat_filtered_split_cropped/test/images \
    -o leucanthemum_flat_filtered_split_cropped_test_prediction \
    -c -v
```

6. apply both models sequentially to new images (here test images for demonstration purposes)

6.1 sliding-window cropping

```
ginjinn utils sw_split \
    -i leucanthemum_flat_filtered_split/test/images/ \
    -o leucanthemum_newdata_sw \
    -s 2056 -p 512
```

6.2 bounding-box detection

```
ginjinn predict leucanthemum_flat_filtered_split_sw_analysis \
    -i leucanthemum_newdata_sw \
    -o leucanthemum_newdata_sw_prediction \
    -c -v
```

6.3 merge sliding-windows

```
ginjinn utils sw_merge \
    -a leucanthemum_newdata_sw_prediction/annotations.json \
    -i leucanthemum_newdata_sw \
    -o leucanthemum_newdata_sw_prediction_merged \
    -t bbox-detection
```

6.4 crop bounding boxes

```
ginjinn utils crop \
    -I leucanthemum_newdata_sw_prediction_merged \
    -o leucanthemum_newdata_sw_prediction_merged_cropped \
    -t bbox \
    -p 25 \
```

6.5 segmentation with refinement

```
ginjinn predict leucanthemum_flat_filtered_split_cropped_analysis \
    -i leucanthemum_newdata_sw_prediction_merged_cropped/images \
    -o leucanthemum_newdata_segmentation_prediction \
    -c -v \
    -r
```
