## Supplementary material for "GinJinn2: Object detection and segmentation for ecology and evolution": seeds.yaml

```
project_dir: /home/ginjinn/seeds_split_analysis
task: "bbox-detection"
# Input data options
input:
  type: "COCO" # or "PVOC"
  training:
    annotation_path: "../seeds_split/train/annotations.json"
    image_path: "../seeds_split/train/images"
  validation:
    annotation_path: "../seeds_split/val/annotations.json"
    image_path: "../seeds_split/val/images"
  test:
    annotation_path: "../seeds_split/test/annotations.json"
    image_path: "../seeds_split/test/images"
# Model options
model:
  name: "faster_rcnn_R_101_FPN_3x"
  weights: "pretrained"

  # additional model options
  model_parameters:
    # anchor generator options
    anchor_generator:
      sizes:
        - - 32
        - - 64
        - - 128
        - - 256
        - - 512
      aspect_ratios:
        - - 0.5
          - 1.0
          - 2.0
      angles:
        - - -90
          - 0
          - 90
    # region proposal network options
    rpn:
      iou_thresholds:
        - 0.3
        - 0.7
      batch_size_per_image: 256
    # regions of interest options
    roi_heads:
      iou_thresholds:
        - 0.5
      batch_size_per_image: 512

# Options for model training
training:
  learning_rate: 0.00125
  batch_size: 1
  max_iter: 7000
  eval_period: 250
  checkpoint_period: 1000
  checkpoint_period: 1000

# Options for image augmentation
augmentation:
  - horizontal_flip:
      probability: 0.25
  - vertical_flip:
      probability: 0.25
  - brightness:
      brightness_min: 0.8
      brightness_max: 1.2
      probability: 0.25
  - contrast:
      contrast_min: 0.8
      contrast_max: 1.2
      probability: 0.25
  - saturation:
      saturation_min: 0.8
      saturation_max: 1.2
      probability: 0.25
  - rotation_range:
      angle_min: -30
      angle_max: 30
      expand: True
      probability: 0.25

# Additional options
options:
  n_threads: 5
  resume: True
```
