## Supplementary material for "GinJinn2: Object detection and segmentation for ecology and evolution": split_image.py

```
import os
import skimage
import skimage.io
import numpy as np

def split_image(img_file, out_dir):
    img = skimage.io.imread(img_file)
    img_gray = skimage.color.rgb2gray(img)
    img_gray = img_gray / 255 if img_gray.max() > 1.0 else img_gray

    img_name = os.path.basename(img_file)
    img_name = os.path.splitext(img_name)[0]

    border_r = img.shape[1] // 10
    stride = 2
    w = 2

    th_m = 0.05
    th_sd = 0.1


    ls = []
    us = []
    ths = []
    for l in range(border_r, img.shape[1] - border_r - w // 2, stride):
        u = l + w
        img_slice = img_gray[:,l:u]
        m, sd = np.mean(img_slice), np.std(img_slice)
        ls .append(l)
        us.append(u)
        ths.append(m < th_m and sd < th_sd)
    ls = np.array(ls)
    us = np.array(us)
    ths = np.array(ths)
    
    if not os.path.exists(out_dir):
        os.mkdir(out_dir)

    if not any(ths):
        print(f'Nothing to do for image "{img_file}"')
        out_file = os.path.join(out_dir, f'{img_name}_x.jpg')
        skimage.io.imsave(out_file, img)
    else:
        img_0_end = ls[ths][0]
        img_1_start = us[ths][-1]

        img_0 = img[:, :img_0_end]
        img_1 = img[:, img_1_start:]

        out_file_0 = os.path.join(out_dir, f'{img_name}_0.jpg')
        out_file_1 = os.path.join(out_dir, f'{img_name}_1.jpg')
        skimage.io.imsave(out_file_0, img_0)
        skimage.io.imsave(out_file_1, img_1)

split_image('PATH TO IMAGE FILE', 'PATH TO OUTPUT DIR')
```
