## Supplementary material for "GinJinn2: Object detection and segmentation for ecology and evolution": stickytraps.yaml

```
project_dir: /home/ginjinn/stickytraps_split_sw_analysis
task: "bbox-detection"
# Input data options
input:
  type: "PVOC" # or "PVOC"
  training:
    annotation_path: "../stickytraps_split_sw/train/annotations"
    image_path: "../stickytraps_split_sw/train/images"
  validation:
    annotation_path: "../stickytraps_split_sw/val/annotations"
    image_path: "../stickytraps_split_sw/val/images"
  test:
    annotation_path: "../stickytraps_split_sw/test/annotations"
    image_path: "../stickytraps_split_sw/test/images"
# Model options
model:
  name: "faster_rcnn_R_101_FPN_3x"
  weights: "pretrained"

# Options for model training
training:
  learning_rate: 0.00125
  batch_size: 2
  max_iter: 5000
  eval_period: 250
  checkpoint_period: 1000

# Options for image augmentation
augmentation:
  - horizontal_flip:
      probability: 0.25
  - vertical_flip:
      probability: 0.25
  - brightness:
      brightness_min: 0.8
      brightness_max: 1.2
      probability: 0.25
  - contrast:
      contrast_min: 0.8
      contrast_max: 1.2
      probability: 0.25
  - saturation:
      saturation_min: 0.8
      saturation_max: 1.2
      probability: 0.25
  - rotation_range:
      angle_min: -30
      angle_max: 30
      expand: True
      probability: 0.25

# Additional options
options:
  n_threads: 5
  resume: True
```
